# The most abundant plant species of Eastern Australia and our knowledge of their traits

**DOI:** 10.64898/2025.12.15.694514

**Authors:** David Coleman, Mark Westoby, Elizabeth Wenk, Laura Williams, Julian Schrader

**Author notes:** Corresponding author: David Coleman.

## Abstract

1. The rule that most species are relatively rare and few are common has been of central importance to many branches of ecology. This rule also implies that just a few species make up the majority of plant biomass in ecosystems. Focussing plant trait research on the most common species in a region could be useful for many applications, including understanding macroscale patterns of species assemblages at the continental scale, identifying strategies of abundant species or estimating landscape-scale fluxes of carbon and water.
2. Here, we rank species based on their total cover across an entire continental region— Eastern and Central Australia—and report the extent of trait coverage in terms of vegetation cover. To calculate species abundance and trait coverage, we used ∼100,000 vegetation plots from the Harmonised Australian Vegetation plot database overlaid across Australia’s vegetation classification scheme (the National Vegetation Information System), and we extracted traits from AusTraits, the most comprehensive regional trait database.
3. Just 113 plant species (<1% of species found in the region) or the combined species of 16 genera make up 50% of the entire vegetation; approximately 10% of species or the combined species from 10% of the genera (1132 species and 140 genera) make up 90% of vegetation cover. Plant trait coverage tended to be higher when expressed as a proportion of vegetation cover than as a proportion of species in the most common woody genera Acacia and Eucalyptus. Gaps in the trait coverage of very common species were obvious, particularly among the grasses of central Australia. Expressing trait coverage weighted by abundance revealed that only a few additional trait measurements of the most common species and genera would be needed to characterise the traits of most of the vegetation in this part of the continent.
4. *Synthesis*: Our results show that a small fraction of species dominates most of the continent. This means that strategic sampling of a few common yet unsampled species could dramatically boost trait coverage and help address the Raunkiaeran shortfall of traits. Targeting these species would substantially improve ecosystem flux estimates and understanding of successful plant strategies at continental scales, with major benefits for vegetation modelling.

## INTRODUCTION

The rule that most species are rare and few are common is a near universal observation in ecology. Ranking species based on their abundance is essential for multiple ecological theories, including understanding plant community assembly processes, how new species evolve and resource partitioning in ecosystems (Grime 1998; Smith *et al*. 2020; Tilman 1990). At larger (continental – global) scales these rankings have been used to prioritise species for conservation, focussing attention on the many rare species (Enquist *et al*. 2019; Fukaya *et al*. 2020). But equally important applications can be found when focussing attention on the few common species (Cooper *et al*. 2024; Zhang *et al*. 2025). In particular, ecosystem function is predominantly determined by the traits of dominant species, as reflected in mass ratio theory (Grime 1998; Smith *et al*. 2020). Given that just 2.2 % of species in tropical forests make up 50% of tree stems in these highly speciose forests (Cooper *et al*. 2024), it is reasonable to assume that other regions composed of a less diverse array of species would display even more extreme skewed abundance distributions. Identifying the most common species in a bioregion, state or continent can be a powerful tool to understanding the organisational structure and function of ecosystems and can serve as a guide for where to focus research efforts to characterise ecosystem function more accurately and efficiently (Ulrich *et al*. 2025).

In order to understand terrestrial ecosystem function, we need good trait coverage across the species that grow there. Traits allows us to group plants by similar function (i.e., plant functional types, PFTs) (Diaz & Cabido 1997; Duckworth *et al*. 2000; Grime 1998; Westoby 2025), model fluxes of carbon and water and identify general patterns of plant functional strategies across environmental gradients. Several traits have emerged as useful for summarising plant function (Diaz & Cabido 1997; Eller *et al*. 2020; Sun *et al*. 2022). These include both categorical traits, e.g., plant growth form and life history, and continuous numeric traits from the plant economic spectrum such as maximum plant height, seed dry mass and leaf mass per area, tissue chemistry traits such as leaf N, leaf P and leaf delta13C, and physiological traits such as turgor loss point (TLP), indicative of drought tolerance or Vcmax, the maximum rate of carboxylation. Such trait data have become readily accessible across species via amalgamation of individual studies into large trait databases (Falster *et al*. 2021; Kattge *et al*. 2020). Identifying gaps in taxonomic trait coverage has become straightforward.

Quantifying trait coverage in this way has greatly accelerated the progress of trait ecology and highlighted the value of complete trait coverage across all species (Wenk *et al*. 2024a). But it has also highlighted the gaps, made conspicuous when performing analyses that require each species included to have observations for the full set of traits. This is one of the many critical challenges to trait based ecology, summarised by the term Raunkiæran shortfalls (de Bello *et al*. 2025; Maitner *et al*. 2023). In these cases and in most trait-based research, all species are treated as of equal importance to the functioning of vegetation communities (Andrew *et al*. 2022; Díaz *et al*. 2022; Towers *et al*. 2024). Statistical gap-filling is frequently employed to carry out trait-environment analysis (Andrew *et al*. 2025; Johnson *et al*. 2021; Joswig *et al*. 2023). The fact that most species are rare and few are common at continental scales means that many of the gaps in trait coverage are likely to be from rare or localised species that contribute comparatively less to ecosystem fluxes of carbon, water and nutrients. If common and widespread species have more trait measurements than rare species with limited distribution, trait coverage expressed in terms of vegetation cover may be substantially greater than trait coverage where all species are given equal weight. Examining trait coverage in terms of the relative abundance of species would also help identify species to prioritise for additional trait measurements.

However, in contrast to *taxonomically* organised trait data – that is how many species have trait data available – assessing the coverage of traits in proportion to their *abundance* – i.e., how much of the vegetation cover has trait available – across the landscape is challenging. There is currently a lack of standardised measures of species abundance at the continental scale, i.e. relative abundance of species across the landscape, rather than within local scale plots. Existing national and state species lists and floras contain no quantitative information of commonness or abundance. Even at local scales, plant survey data tends to be reported in terms of communities or habitats, where species lists are unranked or only partially ranked in terms of abundance. For example, characteristic or dominant plant species may be used to identify vegetation types, but the relative importances of the remaining majority of species are not known (NVIS v.7 2024). Recently, vegetation plot data has been used to derive the number of hyperdominant species in tropical forests using stem counts (Cooper *et al*. 2024) and the relative dominance of species in grasslands using aerial cover and biomass (Zhang *et al*. 2025). Vegetation plot cover data from plots distributed across a region could provide a simple, standardised “best-guess” estimate of species abundance at regional to continental scales. Such a list would enable identification of important gaps in trait coverage and contribute to the understanding of vegetation structure and organisation more generally at the continental scale.

Here we present a list of species ranked by km^2^ of vegetation cover over 4 million km^2^ of Australian continent using the Harmonised Australian Vegetation plot database (HAVplot) (Mokany *et al*. 2022). Specifically we i) rank species in terms of total estimated cover and show the relative abundances of common versus rare species across this part of the continent by growth form and genus, ii) report how the trait coverage in terms of vegetation cover compares with taxonomic trait coverage in the AusTraits plant trait database (Falster *et al*. 2021) and iii) estimate the sampling effort required to fill the gaps in trait coverage for 50% of vegetation cover for this region of Eastern Australia.

## METHODS

### HAVplot database

To estimate vegetation cover of species across Eastern Australia, we used the Harmonised Vegetation plot dataset (Mokany *et al*. 2022), a collation of data from 205,084 vegetation plots sourced primarily from state government agencies and sampled mainly from the 1990’s to 2019. Our study covered the five large states and territories that make up the Eastern Australian mainland: Queensland, New South Wales (including the Australian Capital Territory & Jervis Bay Territory), Victoria, South Australia and the Northern Territory.

Vegetation plot data do exist for the two remaining large states that make up Australia, Tasmania and Western Australia (Mokany *et al*. 2022). However, these were not publicly available at the time of our study. The major steps in data standardisation of HAVplot data, assigning land area to plots and calculating a total cover area per species is described in Figure 1.

**Figure 1.**
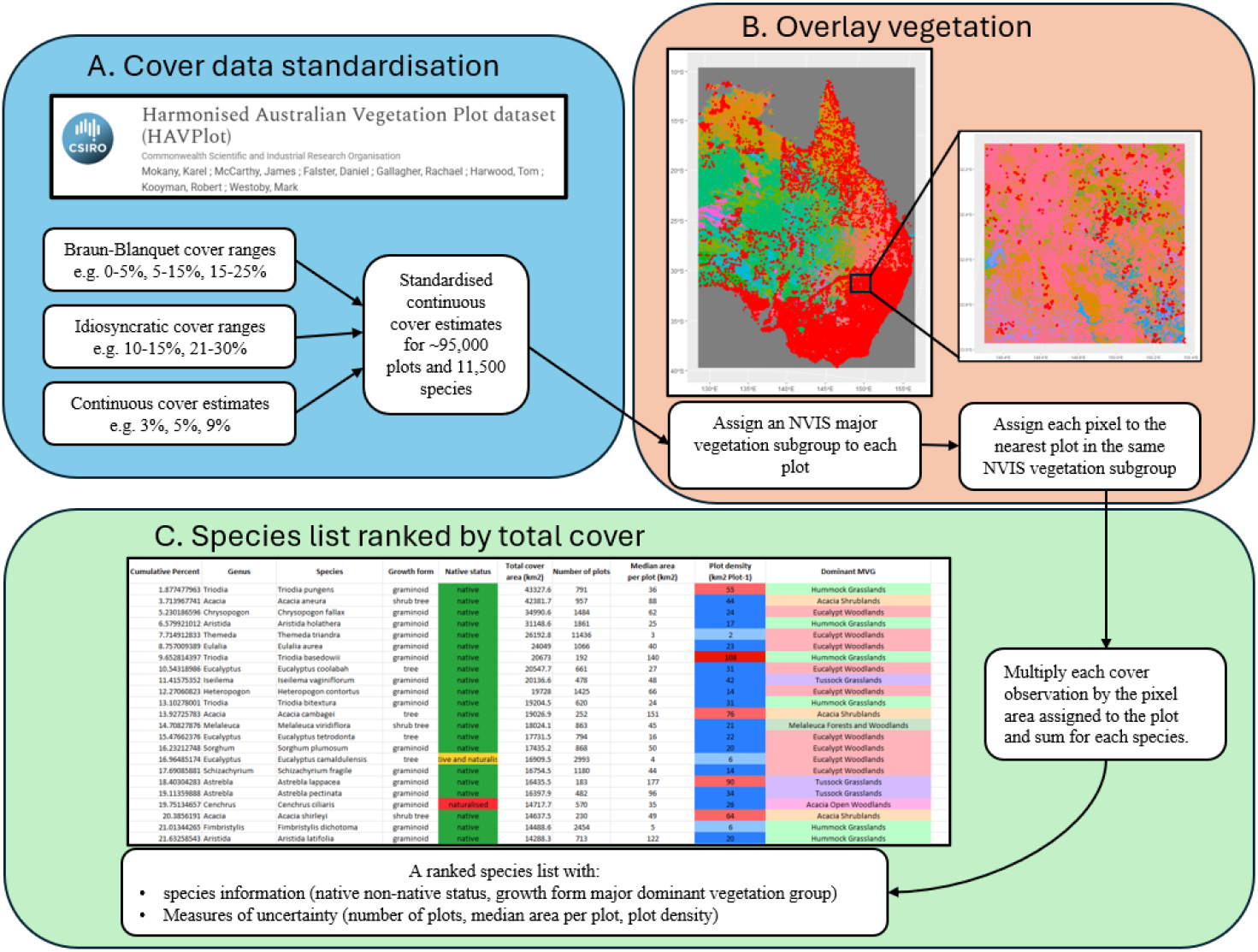
Overview of the methodology to arrive at total cover estimates for each species based on the HAVplot dataset and the NVIS raster map. Red dots in the map extracts in B represent plots and while background colours represent different vegetation classifications in the NVIS scheme. The table cutout in C is a snapshot of Supplementary Data 1 – the list of species ranked by cover area.

### Naming standardisation

A standardised current taxonomic name was attributed to the original species names in this subset using the create_taxonomic_update_lookup function from the APCalign R package (Wenk *et al*. 2024b). A growth form was assigned via a list of standardised growth forms for the Australian flora (Wenk *et al*. 2024a). For taxa that were only identified to genus level this was only possible where all species in a genus had the same standardised growth form. In figures and analyses, non-graminoid herbs, herbaceous climbers and ferns were assigned to the term forbs while species with woody climber, palmoid growth forms and species assigned as both shrubs and trees are assigned to the tree growth form.

### Cover data standardisation

The HAVplot cover estimates required standardisation to a continuous numeric format across the data formats (Figure 1 A). HAVplot vegetation surveys that report vegetation cover estimates contained either ordinal range (2,541,442 cover observations) or continuous numeric (770,190 cover observations) estimates of percent cover. Most of the ordinal values consisted of ranges of percent cover, e.g., 0-5, 5-25, following the Braun Blanquet scale of cover estimation (Braun-Blanquet, 1932), but there were also many other ranges of cover estimate, e.g. 0-1 or 10-15. In any given ordinal category of cover, most species fall towards the lower end of range, creating a skewed distribution of continuous cover estimates. For example, across many species classed in the 5–25% cover category across many plots, the mean continuous cover value might lie close to 9% cover, rather than halfway between (15%). A two-parameter beta distribution is the most widely accepted method for modelling plant cover because it can accommodate asymmetrical distributions constrained between fixed intervals such as the Braun Blanquet distribution categories (Damgaard & Irvine 2019; McNellie *et al*. 2019). To estimate a mean value of continuous cover from the ordinal categorical data, we used the mean estimates of cover published in McNellie et al (2019) based on beta distribution modelling and where possible, tailored to the growth form of the taxon.

The beta distribution consists of two shape parameters which can be fitted to a given expected skewness and the upper and lower boundaries of a curve. For idiosyncratic cover ranges (e.g., 1-4 %, 20-30 %), we inferred a beta distribution using the skewness parameters published by McNellie et al (2019) and we used the mean of this beta distribution as the estimate of continuous cover. In addition to continuous numeric estimates for each cover value, standardised cover ranges based on the Braun-Blanquet scale were assigned to each cover estimate to provide categories of cover for analyses such as in Figure 2. These were 0-5%, 5-25%, 25-50%, 50-75% and 75-100%.

**Figure 2.**
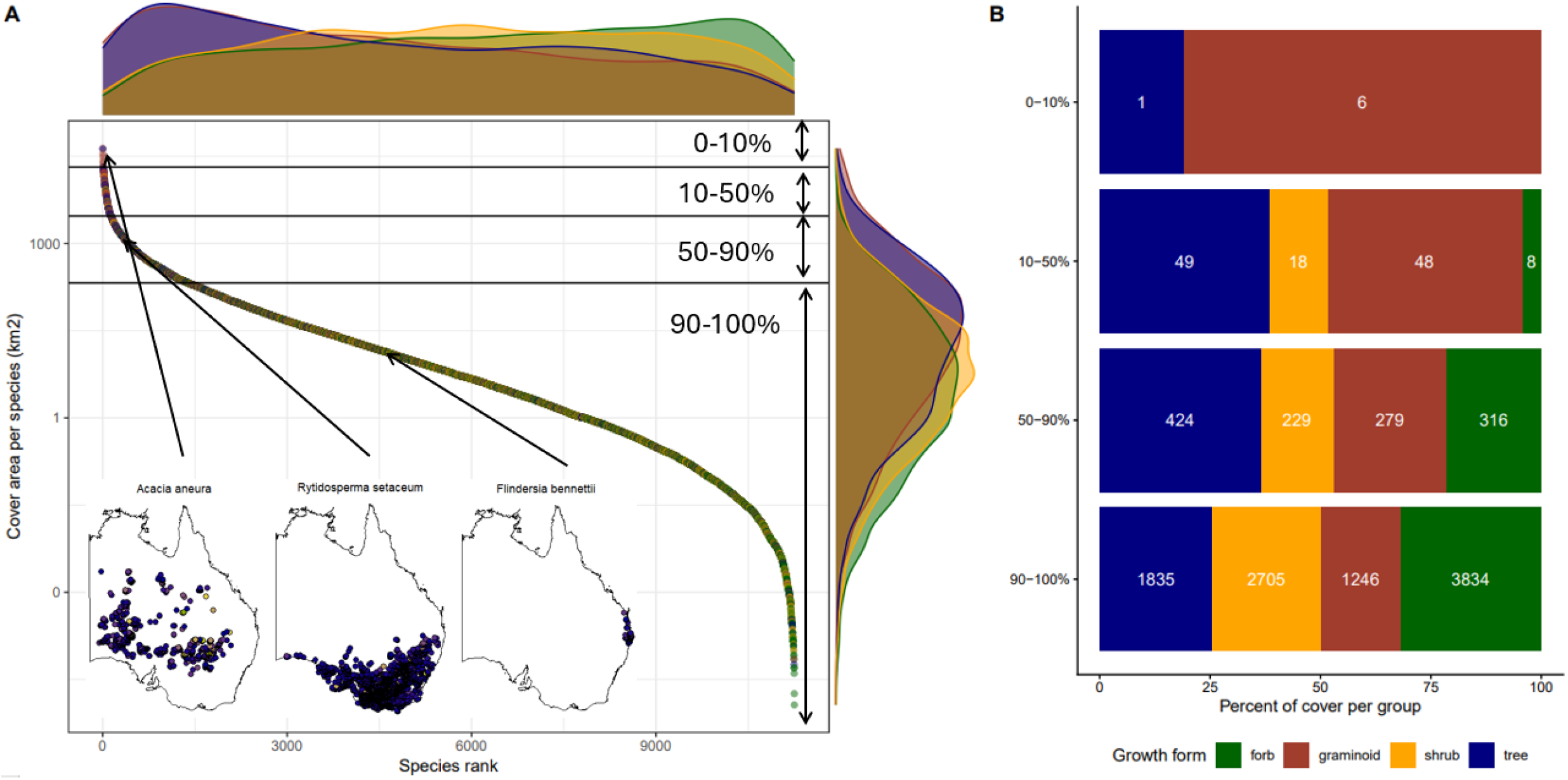
(A) Species ranked by most to least cover area across the region in a log-abundance curve with side panels showing the density of growth forms in both axes. Horizontal lines are at the 10, 50, and 90 cumulative percent total cover area. Maps show the distribution of plots of three example species across the study area and arrows show the approximate position of three example species on the ranked cover scale. Coloured density plots show the relative density of each growth form across both axes of the plot. Note that some rare species at the base of the curve have total estimated covers less than 1 km^2^. These species are likely present in a single plot and not present in nearby adjacent plots. (B) The proportions of growth form in each horizontal band in A. Coloured bands are the proportion of cover per growth form and white numbers are the number of species of each growth form making up the cover. Note that the 7 most common species contribute as much vegetation cover (10%) as >9000 of the least common species. Data underlying the plots in this figure can be found in Supplementary Data 1.

### Data cleaning

A series of other conditions were used to narrow down the vegetation plots to those considered useful for our analysis (Table S1). We removed plots that did not report all vascular species, that did not have a recorded area, that had an area larger than 20 km^2^, where less than 90% of species matched an accepted name in the Australian plant census using APCalign (Wenk *et al*. 2024b). In the few cases where plots shared the same coordinates the plot data with the most recent sampling date at these locations was retained and the rest excluded. We also excluded plots sampled pre 1980 (likely lower accuracy of geographic coordinates), plots without an NVIS vegetation group assigned (outside land area of Australia) and also outside the study area (in Western Australia and Tasmania). This resulted in a list of 96,695 plots used in the analysis.

### Vegetation classification of plots

In order to extrapolate the plot data over the landscape, we assigned major vegetation groups (MVGs) and subgroups (MVS)) to each plot using the National Vegetation Information System (NVIS) version 7, a national map of vegetation types (NVIS Technical Working Group 2017; NVIS v.7 2024) (Figure 1 B). The MVS of the plot coordinates from the vegetation map raster was extracted using the extract() function from the raster package in R (Hijmans *et al*. 2015). The NVIS documentation emphasises that the vegetation type at each 100×100 m pixel in the raster represents the majority inferred from their calculations.

Therefore, some small proportion of plots will be assigned to the wrong vegetation type if the vegetation patch is smaller than the resolution of the map. There are a range of other reasons for vegetation misclassifications using this method, including changes in vegetation cover between plot sampling and the creation of NVIS v7.0, and spatial inaccuracies in the coordinates of the plots and in the pixel classifications of the NVIS map. The NVIS pixel classifications are based on the state vegetation maps. These state vegetation maps are constructed using a variety of methods, including soil maps, remote sensing, expert opinion and modelling, with accuracy varying within regions of state maps (NVIS Technical Working Group 2017). However, NVIS is largely ground-truthed using the vegetation survey data contained in HAVplot and we can thus be reasonably confident that there is good correlation between the two data. As partial validation of the success of the vegetation assignment using NVIS, we looked at the growth forms proportions (Figure S1) across the vegetation classes. In all cases, the dominant growth form corresponded to the description of the vegetation classes, i.e. shrub species dominated shrublands, graminoids dominated tussock grasslands etc.

### Inferring species cover between plots to calculate total species cover

Total cover per species was calculated as a function of the percentage of cover within a plot and the size of the area surrounding the plot within the same vegetation type. First, we classified each plot as belonging to a MVS in the NVIS classification system using the NVIS v7.0 raster. Then each raster cell was assigned to the nearest plot within the same MVS using the knn() function from the FNN package in R (Beygelzimer *et al*. 2015). To reduce memory allocation for these calculations, the NVIS v7.0 raster was broken into 800 smaller tiles (1000 pixels per tile) and the outputs summed. To calculate the amount of total amount of cover per species, the standardised numeric percent cover value from HAVplot was multiplied by the area assigned to each plot and these areas were summed for each species across the landscape within the study area.

Total land area in the study amounted to 5,036,725 km^2^ (shown in Figure S2), about 66% of Australia’s total land area. This includes agricultural land and urban areas and data from these plots were excluded from the analysis. When land area classified as “Unknown/No data” or “Cleared, non-native vegetation, buildings” was removed, this resulted in a reduction of land area of 869,191 km^2^ (∼15%) to the land area in the study. The final land area was 4,167,533 km^2^ (83% of total land area). A total of 19,011 plots (∼20% of plots) were located in either “Unknown/No data” or “Cleared, non-native vegetation, buildings” vegetation classification zones (Table S2).

All vegetation cover estimates for species are expressed in terms of vegetation cover area, not in terms of geographic land area. Total cover in each plot did not necessarily sum to 100%. In dense vegetation with many strata, total cover in the plot would sum to well over 100%. On the other hand, many plots in the large arid interior of Australia had only a fraction of the land area covered by vegetation (i.e. the area would have a leaf area index considerably less than 1), the area of total vegetation cover in this part of the continent was less than the land area over which the cover was spread. There was 2,381,494 km^2^ of vegetation cover spread (or stacked in three-dimensional space, for example in forested plots with multiple strata) across the 4,167,533 km^2^ land area (totalling 57% of the land area), consisting of 2,666,159 species observations. 85,038 (∼3%) of these species-by-cover observations could not be assigned to an accepted species name in the APC or were identified to the genus level and were excluded from the analysis. This resulted in a further reduction in total vegetation cover of 151,199 km^2^ to 2,289,533 km^2^. All further analyses were based on this final list of 11,251 species and their inferred total area 2,289,533 km^2^.

The results for this process are summarised in Supplementary Data 1 and 2, ranked lists of species and genera by total cover area. Each species is accompanied by growth form, native/non-native status and the dominant MVG from the NVIS scheme. As a measure of confidence in the estimated total cover area and ranking, the number of lots, median area assigned to each plot and the plot density (total area divided by the number of plot observations) accompanies each species.

### Method validation

We assigned vegetation types to each plot using the NVIS to be as representative as possible in extrapolating vegetation cover to the appropriate landscape surrounding them. For example, two plots may be positioned each side of a boundary between vegetation types and one of them is much further from the boundary than the other. Without incorporating NVIS in the calculation, the area between them would be divided equally between the two plots. We calculated an alternative estimate of total cover and species rankings without using the NVIS and compared the rankings of species between the two methods (Figure S3). We calculated total cover per species using Voronoi panels; that is, where the land area was divided into regions based on the proximity to the nearest plot, rather than the nearest plot within the same vegetation type. This process was carried out using the deldir R package (Turner & Turner 2015).

In order to assess the completeness of the flora list from these ∼95,000 plots we also scrutinised the list of species in our study to see how many of the species that are officially recognised to occupy the study area by the Australian Plant Census (APC 2025) were captured in our analysis and importantly, how many were not. A complete list of Native and Non-native species known to inhabit the states of NSW, Queensland, Victoria, Northern Territory and South Australia was assembled from the APC and compared to our list of species with cover data in HAVplot. 67% of the estimated 16,874 species that are present according the Australian Plant Census (APC 2025) exist in the study. To see how the ranking and cover estimates for species compared with numbers of occurrence records in herbaria, we used the galah R package (Westgate *et al*. 2024) to download total occurrence record counts in the Australian Virtual Herbarium (AVH 2025) for each species as a measure of whether we captured common species as would be defined by total number of herbarium records.

### Trait coverage

Species coverage for 11 representative traits used in characterising plant functional types and ecosystem function models were extracted from the AusTraits plant trait database v 7.0.0 (Falster *et al*. 2021) using the extract_trait function in the austraits R package. These traits were chosen because they represent the plant economic spectrum (Díaz *et al*. 2016) and are commonly used in ecosystem models (Marshall *et al*. 2008). The traits we selected are as follows: Plant height, Seed dry mass, Leaf Mass per Area (LMA), Leaf Nitrogen per dry mass (Leaf N), Leaf Phosphorous per dry mass (Leaf P), Leaf area, Leaf delta13C, Leaf Turgor Loss Point (TLP), Leaf photosynthesis Vcmax per unit area (Vcmax), Leaf stomatal conductance per area at Amax (g_smax_) and Wood density. Definitions of these traits can be found in the AusTraits plant trait dictionary (Wenk *et al*. 2024c). We considered a species to have trait coverage if at least one trait value was present in the AusTraits database for that trait and species. We combined this trait information with the landscape cover estimates above to produce an estimate of trait coverage of the vegetation in the study area of Eastern Australia grouped by genus (results shown in Table 1) and also species ranked by increasing cover (results Figure 2). Grouping by genera as well as by species is useful for two reasons: i) splitting taxa into species can be contentious, particularly across morphologically similar taxa such as Eucalypts and grasses and ii) during data collection, vegetation surveyors could unintentionally attribute similar looking but rarer species within a genus to the more common one.

**Table 1.** The 16 genera that make up 50% of the vegetation cover across Eastern Australia and the proportion (between 0 and 100%) of this vegetation cover that has trait data. Traits were selected to represent a range commonly used in plant trait ecology. Coloured squares show increases or decreases in trait coverage using proportion of vegetation cover compared with proportion of species. Positive changes (green colour) represent greater coverage when using proportion of vegetation cover. NA values for the wood density trait are for herbaceous genera. Trait names LMA and gs max represent Leaf Mass per Area and Maximum rate of stomatal conductance respectively. C3/C4 refers to the Photosynthetic pathway of species in each genus. Data underlying this table can be found in Supplementary Data 2

| Family | Genus | n species | Growth form | C3/C4 | Total Cover (km2) | n observations | Cumulative percent cover | Plant height | Seed mass | Leaf area | LMA | Leaf N | Leaf P | Leaf delta13C | TLP | gs max | Vcmax | Wood density |
| --- | --- | --- | --- | --- | --- | --- | --- | --- | --- | --- | --- | --- | --- | --- | --- | --- | --- | --- |
| Myrtaceae | Eucalyptus | 352 | tree | C3 | 318179 | 127964 | 13.79 | 99 | 95 | 87 | 93 | 88 | 57 | 84 | 35 | 31 | 21 | 86 |
| Fabaceae | Acacia | 459 | tree | C3 | 205164 | 80277 | 22.68 | 100 | 94 | 69 | 72 | 65 | 51 | 67 | 22 | 22 | 0 | 72 |
| Poaceae | Triodia | 29 | graminoid | C4 | 132920 | 4246 | 28.45 | 100 | 83 | 68 | 11 | 26 | 26 | 6 | 0 | 3 | 0 | NA |
| Poaceae | Aristida | 49 | graminoid | C4 | 87400 | 20660 | 32.23 | 100 | 61 | 99 | 73 | 54 | 54 | 38 | 0 | 0 | 0 | NA |
| Myrtaceae | Melaleuca | 68 | tree | C3 | 47896 | 11317 | 34.31 | 91 | 87 | 71 | 77 | 73 | 64 | 70 | 5 | 1 | 0 | 80 |
| Poaceae | Astrebla | 4 | graminoid | C4 | 44867 | 1141 | 36.26 | 100 | 100 | 100 | 73 | 27 | 27 | 0 | 0 | 0 | 0 | NA |
| Poaceae | Eragrostis | 68 | graminoid | C4 | 43555 | 12368 | 38.14 | 96 | 91 | 80 | 51 | 2 | 2 | 4 | 0 | 0 | 0 | NA |
| Poaceae | Chrysopogon | 10 | graminoid | C4 | 40653 | 2048 | 39.91 | 100 | 89 | 91 | 0 | 86 | 86 | 86 | 0 | 0 | 0 | NA |
| Amaranthaceae | Sclerolaena | 47 | herb/shrub | C3 | 35415 | 14152 | 41.44 | 100 | 84 | 87 | 77 | 17 | 0 | 17 | 0 | 0 | 0 | 0 |
| Scrophulariaceae | Eremophila | 50 | shrub | C3 | 32667 | 8231 | 42.86 | 100 | 95 | 84 | 86 | 83 | 63 | 82 | 35 | 31 | 0 | 48 |
| Poaceae | Themeda | 4 | graminoid | C4 | 28660 | 11721 | 44.10 | 100 | 96 | 100 | 91 | 91 | 91 | 91 | 0 | 0 | 0 | NA |
| Amaranthaceae | Atriplex | 40 | shrub | C4 | 28102 | 7708 | 45.32 | 100 | 96 | 90 | 87 | 22 | 22 | 9 | 0 | 15 | 0 | 16 |
| Amaranthaceae | Maireana | 40 | herb/shrub | C3 | 28085 | 10858 | 46.53 | 100 | 98 | 82 | 76 | 58 | 24 | 34 | 0 | 24 | 0 | 0 |
| Poaceae | Heteropogon | 2 | graminoid | C4 | 27205 | 2283 | 47.71 | 100 | 100 | 100 | 27 | 100 | 73 | 73 | 0 | 0 | 0 | NA |
| Poaceae | Enneapogon | 15 | graminoid | C4 | 26892 | 3339 | 48.88 | 100 | 100 | 85 | 80 | 26 | 26 | 0 | 0 | 0 | 0 | NA |
| Poaceae | Sorghum | 12 | graminoid | C4 | 26874 | 2700 | 50.04 | 100 | 25 | 85 | 0 | 14 | 14 | 0 | 0 | 0 | 0 | NA |

Change in trait coverage

To test for significant differences in areas covered among the major growth forms (i.e., forbs, graminoids, shrubs, trees) we conducted an ANOVA on the log_10_ areas of species, grouped by growth form. The aov() and TukeyHSD() functions in base R were used to compute significant differences and the 95% CI for the means of each group.

All data wrangling and plots were carried in Rstudio (Posit team 2025) dplyr package (Wickham *et al*. 2023) and using base R functions and the ggplot2 package (Wickham 2016) and the code for repeating the analysis can be found at https://doi.org/10.6084/m9.figshare.30889301 Figshare repository.

## RESULTS

We found that just 132 species or 1.2% of the 11,122 species observed in the plots across our study area contributed 50% of estimated vegetation cover (Figure 2 A and B, Supplementary Data 1) across eastern continental Australia, a region spanning deserts, grassy woodlands, tall open forests and rainforests (Figure S4) (NVIS v.7 2024). The skewed distribution is such that just seven common species (six grasses and one tree species) were estimated to cover the same area as >9000 of the least common species. This same pattern of highly skewed species distributions was found when species were grouped into growth forms of trees, forbs, shrubs and graminoids (Table S3), despite forbs contributing far less to the top 50% of vegetation cover than would be anticipated by their species richness (Figure 2 B). The most common species were mainly trees and graminoids. The median cover area per species for trees and graminoids (12.9 and 14.6 km^2^ respectively) were statistically similar and 3-6 times larger (ANOVA on log10 areas in km^2^ df = 3, F = 197.3, p < 0.001) than for shrubs and forbs (4.0 and 2.5 km^2^). The cover area per species correlated linearly with number of vouchers in the AVH (n: 11,251, p < 0.001, R^2^: 0.37; Figure S4 and Table S3).

Vegetation cover of common genera across the continent was taxonomically narrow. Just 16 genera (< 1% of all genera) made up 50% of vegetation (Table 1). Of this 50%, woody genera *Eucalyptus, Acacia* and *Melaleuca* accounted for ∼25% of vegetation cover with the remaining 25% composed mostly of C4 grasses and shrubs.

For these common genera, trait coverage tended to be greater in terms of vegetation cover than in terms of the fraction of species (green squares, Table 1). Genera containing tree species tended to have trait information for more than half their vegetation cover, with the exception of the physiological traits TLP, g_smax_, and V_cmax_. In contrast, common species in some Poaceae genera (e.g. *Triodia, Eragrostis, Sorghum*) were undersampled, even in terms of traits such as LMA and leaf N that had high coverage in other prominent genera. If sampling efforts were targeted at the most common 1% species, we estimate at most ∼120 species per trait are required to reach trait coverage of 50% of vegetation in this part of the continent (Table 2).

**Table 2.** The number of species required to be measured to have complete trait coverage for the most common plant species (i.e., the 132 species that are the top 1% of species and account for 50% of total vegetation cover) by cover for 10 representative traits in the AusTraits database. The final column represents the median percent gain in trait coverage for each additional species measured from among these missing common species.

| Trait name | Number of species to complete the top 50% of vegetation cover | Current proportion of vegetation cover | Percent gain in coverage per species measured |
| --- | --- | --- | --- |
| Plant height | 0 | 97.7 | - |
| Leaf mass per area | 36 | 61.2 | 0.42 |
| Seed dry mass | 12 | 85.7 | 0.37 |
| Leaf N per dry mass | 49 | 49.1 | 0.36 |
| Leaf P per dry mass | 72 | 35.1 | 0.34 |
| Leaf delta13C | 68 | 40.4 | 0.36 |
| Leaf area | 19 | 75.3 | 0.30 |
| Leaf turgor loss point | 115 | 10.5 | 0.36 |
| Leaf photosynthesis Vcmax per area | 124 | 3.2 | 0.38 |
| Leaf stomatal conductance per area at Amax | 117 | 7.6 | 0.37 |

## DISCUSSION

Our study revealed that vegetation cover across eastern continental Australia is dominated by a small subset of species, with cover from only six grasses and one tree species equivalent to more than 9,000 of the least common species. Just 16 genera account for 50% of vegetation highlighting the potential for simplifying the diverse flora into plant functional types.

Expressing trait coverage as a proportion of total vegetation cover highlights that we know more about the vegetation at the continental scale than is suggested by taxonomic coverage alone. It also highlights obvious gaps in the trait coverage of species that make up large parts of the vegetation and a tractable list of species to target for future trait sampling, providing clear guidelines to address the Raunkiaeran shortfall of trait data gaps across scales.

We found the combined flora of Eastern Australia, an area dominated by semi-arid vegetation, to exhibit an abundance distribution that was more highly skewed than of tropical forests across the globe; 1.2 % of species comprised 50% of vegetation cover in comparison to the 2.2 % of species contributing 50% of tree stems > 10cm DBH reported in Cooper et al (2024). This was driven by a common species assemblage largely composed of trees and graminoids which had species cover areas that were an order of magnitude greater on average than shrubs and forbs (Figure 2). Despite the uneven contribution of growth forms to total cover, the highly skewed distribution in terms of cover was also consistent *within* different growth forms: 2.2 % of tree species made up 50% of tree cover and 2.4 % of forbs made up half of all forb cover (Table S3). Furthermore, the skewness of cover area among forbs, the most diverse grouping of species closely resembled that of tree stems in tropical forests (Table S3 and Extended Data Table 5 from Cooper *et al* (2024)). This suggests underlying patterns in commonness and rarity are highly conserved not only within a single growth form and within a single biome but also across a range of metapopulations, functional groupings and biomes at large geographic scales.

Trait coverage in terms of vegetation cover among the most common species revealed that we know more about the vegetation in this part of the continent than would be suggested by the trait coverage by fraction of species alone (Table 1). This was especially true for the three dominant woody genera: Eucalyptus, Acacia and Melaleuca, where even infrequently measured traits like Turgor Loss Point (TLP) or g_smax_ (Maximum stomatal conductance) improved when expressed as a proportion of total cover (green colours) instead of species richness. Among species in common grass and shrub genera, there were gaps in the trait coverage of common species, indicated by reductions (pink colours) or 0’s in the estimate of trait coverage, e.g. Leaf N in *Astrebla* or Seed mass and LMA in *Sorghum*. This is likely due to species selection in research studies. These grasses are abundant across vast areas of sparsely populated parts of the continent. Practicality dictates species selection in ecological and ecophysiological studies be based on what is available in nurseries or seed suppliers, what is possible to cultivate or what is easily accessible in field studies (e.g. common garden close to research facilities, National Parks within a days drive). Abundant species in more isolated regions are therefore more likely to be neglected. Even relatively common species such as grasses that grow close to population centres may be overlooked due to convenience of form or anatomy for trait measurement or a lack of “charisma” (Mesaglio et al. 2023).

Encouragingly, this study provides some quantitative evidence for refocussing attention to such neglected species groups; each new species that is measured in the top 132 species will provide an average increase of 0.3 – 0.4 % trait coverage of the entire vegetation cover in this part of the continent (Table 2).

To our knowledge, this is the first study to rank the majority of a flora in terms of abundance at such a large scale and estimate trait coverage in this way. This means we have little comparison with other regions save for studies focussing on subsets of floras such as rare species or woody species (Fukaya *et al*. 2020; Yilangai *et al*. 2023). Currently, trait coverage in major trait databases of the ∼ 354,000 plant species known to science is low for most traits (for example 12% of species globally for seed dry mass (Díaz *et al*. 2022)). It seems likely that global trait coverage would improve for all traits when expressed in terms of total vegetation cover, given the ubiquitous nature of the few common and many rare rule of species assemblages across continents (Cooper *et al*. 2024). These results provide baseline statistics (e.g. median graminoid total cover area in this study being 14.6 km^2^) for the extent and organisation of vegetation over a largely flat and arid landmass. This could be compared to similar arid or contrastingly more productive landscapes on other continents, particularly in Europe and North America with comprehensive vegetation plot coverage of the landscape (Bruelheide *et al*. 2019; Sporbert *et al*. 2020).

In previous efforts to characterise vegetation function and fluxes of carbon and water, the vegetation community has been simplified by dividing the vegetation up into PFTs (Duckworth *et al*. 2000). It is also possible to quickly attribute the species in the common genera in this study (Table 1) into rudimentary PFTs based on their growth form and photosynthetic pathway; many of the most common species appear to be closely related and are functionally similar e.g. 10 of the 16 genera containing C4 grasses.

These species are those most abundant at the continental scale. But further scrutiny of the abundance rankings organised by genera and growth forms across the flora at more local scales may reveal equally clear patterns with which to construct PFTs (Examples in Figure S5). 90% of the vegetation cover in these biomes are composed of just a fraction (10-20% in Figure S5) of total species which would reduce the number of species to target in order to achieve complete trait coverage and characterise ecosystem function. Furthermore, common species are likely to be present and abundant in multiple communities, reducing even further the gaps in trait coverage.

There are some caveats to using cover area as a proxy for the importance of species to ecosystem functioning. Firstly, vegetation that covers the most area in Australia may be exchanging much less water and carbon than the more productive southern and coastal regions with higher species richness (Figure S6). Dominant, perennial species in the more productive environments (many of which may still found in the top 1330 most common species corresponding to 90% of vegetation cover) could be just as important in terms of total ecosystem process as some of the species with conservative or ephemeral strategies that cover vast areas of the arid zone. Secondly, cover is two dimensional for a given species and does not reflect the three-dimensional structure of height, canopy height and LAI for individual species. Possible extensions of the cover ranked data in this study might be to estimate conversion factors between total cover area and biomass or ecosystem flux (Pan *et al*. 2024). New rankings in terms of ecological or economic importance could then be applied, such as ranking by carbon storage. Thirdly, our estimates of trait coverage by vegetation cover are optimistic. Capturing intraspecific variation of traits in response to the environment is often required to accurately capture ecosystem fluxes (Marshall *et al*. 2008; Sun *et al*. 2024; Zeng *et al*. 2020). While we have reported trait coverage as the bare minimum – at least one datapoint per species – this study provides a starting point for future targeting of species over physiological ranges and conditions that they will experience under climate change projections.

We would like to emphasise that these estimates of cover area should be taken as a guide of commonness and rarity at the continental scale and not as final numbers for individual species. An alternate calculation method produced minimal change for most common species (Residual se: 2 km^2^, Figure S4) but a few outlier species changed their rankings by more than 2000 places. In particular we estimated that our HAVplot dataset missed ∼ 5000 plant species that did not appear in any plot with abundance measures. However we know these missed species are already the rarest in terms of occurrence data (Table S4) and unlikely to change the ratio of common to rare species in terms of cover area; we interpret these species as being below the veil line as part of any species abundance distribution based on sampled data (Preston 1948).

## Acknowledgements

The authors wish to thank Karel Mokany for his assistance compiling the HAVplot dataset. We also thank the many botanists and researchers who visited the plots and sampled the sampling vegetation and trait data used in HAVplot and AusTraits databases. We thank Daniel Falster for building the AusTraits database, his role in the HAVplot database and building the APCalign R package. DC was supported by the Genes to Geosciences foundation.

## CONCLUSION

This first quantification of species abundance across the entire flora of more than 4 million km^2^ of Eastern Australia reveals a highly skewed distribution of species cover areas (Supplementary Data 1). This pattern was consistent when grouping species by growth form and highlights the consistency of such patterns in species abundance assemblages across floras, regardless of biome or region. We found that estimates of trait coverage improved when weighting species by their cover across the landscape – we know more than we thought and this fact is likely to apply on other continents and also globally. Ranking species by cover can be useful for quickly assigning most of the vegetation cover in the region to common PFTs and to identify and encourage the study of less charismatic or convenient but highly abundant species – particularly when there are relatively few in the context of the entire flora. Such a ranked list could serve as a starting point for further projects quantifying landscape level fluxes of carbon and water by conversion of plant cover into biomass and for understanding patterns in vegetation more generally at the continental scale in Eastern Australia.

